# Awareness of danger inside the egg? Evidence of innate and learned predator recognition in cuttlefish embryo

**DOI:** 10.1101/508853

**Authors:** Nawel Mezrai, Lorenzo Arduini, Ludovic Dickel, Chuan-Chin Chiao, Anne-Sophie Darmaillacq

## Abstract

Predation exerts one of the greatest selective pressures on prey organisms. Many studies showed the existence of innate anti-predator responses mostly in early stages of juvenile’s vertebrate. Learning is also possible but risky since it can cause death. There is now growing evidence that embryonic learning exists in animals but few studies have tested explicitly for learning in embryos. Here, *Sepia pharaonis* cuttlefish embryos respond to the presence or odour of a predator fish but not to a non-predator fish. Interestingly, embryos can learn to associate a non-threatening stimulus with an alarm signal: cuttlefish ink. After several paired exposures, they respond to the harmless fish as if it were dangerous. Our results demonstrate both innate and acquired predator recognition in a cephalopod. Embryos response is a decrease of ventilation rate. Such response is adaptive, especially in a translucent egg, since it reduces movement and hence the risk of being detected; this freezing-like behaviour may also reduce bioelectric field, which lessens shark predation risk. Last, our result is the first report of associative learning in an invertebrate embryo. This shows that a cuttlefish embryo can have both genetic predator avoidance skills and possesses enough cognitive abilities plasticity to learn and retain new threats before hatching. The combination of these mechanisms is an impressive example of the early adaptability of cephalopod molluscs. This amount of behavioural plasticity gives the newly-hatched sepia a huge selective advantage when dealing with known or new threats.

## Introduction

From the first moments of life, an individual must be able to protect itself from predators and find food. Young must recognize predators very early in order to avoid them and survive. Predator recognition has a strong innate component. In mammals, birds, amphibians, reptiles, fishes or snails, preys use chemical, visual or auditory cues to recognize predators (Amo, López, & Martín, 2005; Balderas-Valdivia, Ramírez-Bautista, & Carpenter, 2005; Barreto, Luchiari, & Marcondes, 2003; M. M. Brown, Kreiter, Maple, & Sinnott, 1992; Dalesman, Rundle, Coleman, & Cotton, 2006; Dalesman, Rundle, & Cotton, 2007; Fendt, 2006; Griffiths, Schley, Sharp, Dennis, & Roman, 1998; Hartman & Abrahams, 2000; Hawkins, Magurran, & Armstrong, 2004; Hirsch & Bolles, 1980; Saunders, Ong, & Cuthbert, 2013). Some preys, however, require learning to respond to predation. Learned predator recognition has been shown in a diverse range of taxa: birds (Curio, Ernst, & Vieth, 1978); mammals (Kindermann, Siemers, & Fendt, 2009); fishes (Chivers & Smith, 1998; Kelley & Magurran, 2003; Mathis & Smith, 1993); amphibians (Chivers & Smith, 1998; Epp & Gabor, 2008; Ferrari, Manek, & Chivers, 2010; Mathis, Ferrari, Windel, Messier, & Chivers, 2008; Mirza, Ferrari, Kiesecker, & Chivers, 2006; Wisenden, 2003; Woody & Mathis, 1998) and invertebrates (Aizaki & Yusa, 2010; Ferrari, Messier, & Chivers, 2008; Rochette, Arsenault, Justome, & Himmelman, 1998; Wisenden, 2003; Wisenden, Chivers, & Smith, 1997; Wisenden & Millard, 2001). One mode of learning is through the pairing of cues from a predator with an alarm cue (classical conditioning). Indeed, in marine environment, usual way for prey to detect and identify predators are olfactory and visual cues (G. E. Brown & Smith, 1998; Hartman & Abrahams, 2000; Kats & Dill, 1998; Miklósi, Pongrácz, & Csányi, 1997; Utne-Palm, 2001).

Predator recognition can be learned early in development. While protected in egg, embryos can perceive environmental stimuli to identify risk factors that may be present in their post-hatching environment. This embryonic learning ability has been highly studied in amphibians (Ferrari & Chivers, 2009a, 2009b, 2010; Ferrari, Crane, & Chivers, 2016; Ferrari et al., 2010; Golub, 2013; Mathis et al., 2008; Saglio & Mandrillon, 2006). The first study explicitly showing these abilities to recognize predators was conducted by Mathis et al. (2008). It showed that when salamander eggs (*Ambystoma annulatum*) were exposed to chemical predatory cues, larvae showed anti-predatory behaviors such as shelter seeking and reduced locomotor activity (Mathis et al., 2008). Subsequently, further studies have shown that predator recognition can also be learned and generalized to other similar predators (Ferrari & Chivers, 2009b). By observing post-hatching responses, Ferrari and colleagues have shown that amphibian embryos can learn to chemical cues before hatching by using associative learning: predator cues, cues about their diet and/or alarm signal such as smell of congeners injured (Ferrari & Chivers, 2009b, 2009a, 2010; Ferrari et al., 2010; Garcia, Urbina, Bredeweg, & Ferrari, 2017).

Cuttlefish are oviparous Cephalopod Molluscs. Embryos develop in a soft egg elastic case and juveniles do not receive direct parental care because adult die soon after mating (males) or egg-laying (females)(Peter R. Boyle, 1987; Lee, Lin, Chiao, & Lu, 2016)., Romagny and collaborators (2012) showed in *Sepia officinalis* that the different sensory systems are functional before hatching: they observed mantle contractions after tactile, olfactory and light stimulations. Furthermore, other studies highlighted indirect prenatal learning (Darmaillacq, Lesimple, & Dickel, 2008; Darmaillacq, Mezrai, O’Brien, & Dickel, 2017; Guibé, Poirel, Houdé, & Dickel, 2012). Indeed, cuttlefish embryos exposed to small crabs before hatching prefer crabs rather than their innately preferred shrimp prey (Darmaillacq et al., 2008). Likewise, cuttlefish that innately preferred black crabs will preferentially select white crabs following embryonic exposure to them (Guibé et al., 2012). Unlike *Sepia officinalis* in which the egg case is darkened by the maternal ink, in the pharaoh cuttlefish (*Sepia pharaonis*) eggs are totally transparent. This allows a direct observation of the embryos’ responses to both external chemosensory and visual stimuli and thus embryonic learning abilities without modifying the egg capsule.

The aim of this study is to examine if *Sepia pharaonis* embryos can recognize predators innately or if they have to learn to recognize them. To test their innate visual and chemical recognition capabilities embryos will be exposed to predatory and non-predatory cues. To test their learned visual and chemical recognition capabilities, a classical conditioning procedure will be used by pairing a neutral stimulus (the sight or the odour of a non-predatory fish) with an alarm signal: cuttlefish ink. Indeed, the cuttlefish ink can be a relevant warning signal (Derby, 2014). It is composed of secretions from two glands: (1) the gland of the ink bag that produces a black ink containing melanin; (2) the gland in the funnel that produces mucus. The cuttlefish ink is composed of melanin but also catecholamines, DOPA and dopamine (which are monoamines derived from tyrosine), amino acids such as taurine but also certain metals such as cadmium, copper and lead (Derby, 2014; Madaras, Gerber, Peddie, & Kokkinn, 2010; Prota et al., 1981). The ink of cephalopods would then have a role of defense against predators in two ways: (1) ink as a direct deterrent of predators (interspecific effects) and (2) Ink as an alarm cue for conspecifics (intraspecific effect) (Derby, 2014; Hanlon & Messenger, 2018). This second type of defense acts indirectly against predators, because it signals a danger to conspecifics. We hypothesize that embryos can use chemical cues to recognize predator like many vertebrate and some invertebrates (i.e. amphibians) species but also visual cues because of the characteristics of the egg case. Likewise, we hypothesize that embryos can learn about a new danger through chemosensory and visual cues by associative learning. Unlike Romagny et al (2012), ventilation rate (VR) was used as a behavioural measure rather than mantle contractions because VR can be used to monitor more subtle responses to low intensity stimuli (Boal & Ni, 1996). Indeed, in addition to mantle contractions, decreased ventilation and bradycardia can be observed in cuttlefish after sudden visual or chemical stimulation (King & Adamo, 2006). Unlike heart rate, VR is easily and directly observable in cuttlefish and can be easily observed under a microscope, either by noting the rhythmic motion of the collar flaps circulating oxygenated water to the gills, or by the movement of the funnel in response to pressure changes resulting from respiratory movements (inhalation and exhalation).

## Materials and Methods

### 1) Biological model used

#### Experimental model

Model species is the pharaoh cuttlefish (*Sepia pharaonis*). The pharaoh cuttlefish is one of the most important fishery species of cephalopod and is widely distributed from the east Africa to the west Pacific Ocean (Anderson et al., 2010). Adults (4 females and 2 males) were fished and reared in a semi-natural area in Academia Sinica Marine Research Station or Aquaticlch Biotech Company Ltd. aquaculture (Yilan, Taiwan). All the eggs studied were laid in the same location (first generation) and were transferred before organogenesis to the Institute of Systems Neuroscience & Department of Life Science (National Tsing Hua University, Taiwan). Transfer was done in large containers (30×50×30cm) fill with natural seawater. A bubbler pump was installed to give oxygen in the containers. In the institute, eggs were maintained in natural sea water with constant renewal at 25 ± 2°C temperature and on a 12:12 h light:dark cycle. Each egg was separated individually from the clusters and incubated in a plastic basket floating in the culture tanks (20 eggs maximum per basket of 15×20×3cm). The volume of each tank was 300L.

#### Embryonic development

The time course of the development for each egg is different because the eggs are laid singly, and the spawning periods may last for several days. It took 22-24 days to complete embryonic development at a water temperature between 22-25°C (M.-F. Lee et al., 2016). Based on morphological characteristics, 30 stages were observed during the embryonic development of *S. pharaonis:* cleavage from stage 1 to stage 9; blastulation and gastrulation from stage 10 to stage 15 and organogenesis from stage 16 to stage 30 (M.-F. Lee et al., 2016). During the embryogenesis the sensory systems start to develop and become functional (Romagny, Darmaillacq, Guibé, Bellanger, & Dickel, 2012) and embryos respond and recognize chemicals cues (Mezrai, Chiao, Dickel, & Darmaillacq, in preparation). Indeed, embryos respond to predator odour (Narrow-lined puffers: *Arothron manilensis*) but not to non-predator odour (Clownfish: *Amphiprion percula*) from stage 25 embryos (Mezrai et al., in preparation).

#### Chemical and visual stimuli

Different chemical or visual stimulations were presented to cuttlefish embryos: predatory fish, non-predatory fish and cuttlefish ink.

1) Fishes: the predators used were the narrow-lined puffers (*Arothron manilensis*). Two groups of puffers were used: the first group was fed daily on standard food (defrosted shrimp). The second group was given daily one cuttlefish egg. The non-predators, clownfishes (*Amphiprion percula*), were fed *ad libitum* on standard herbivorous aquarium food. All fishes had comparable sizes (4 to 6 cm) and similar swimming activity in the experimental tank (size: 20×60×30cm).

2) Ink: ink was obtained by stressing one-week-old cuttlefish placed in a 300 ml glass container by approaching a net until the container got saturated with ink (i.e. the water was totally black and the cuttlefish were not visible); the cuttlefish were then returned to their hometank.

This procedure was repeated each day. All fishes and cuttlefish used were in natural seawater (25 ± 2°C) with constant renewal and sufficient oxygenation (bubbler pump installed in each aquarium) and on a 12:12 h light:dark cycle.

### 2) Protocol and experimental apparatus

All experiments were conducted in a 36×22×25cm tank totally opaque in order to isolate the experiment from any external visual interference (cf. figure 1). Embryonic behaviour was recorded with an underwater camera (Olympus Stylus Tough TG-4). The cuttlefish egg (stage 25) was placed on the bottom in the center of the tank for 5 min (acclimation phase) on a plastic stand to prevent it from rolling. Then, the olfactory or the visual stimuli were presented to the embryo (stimulation phase).

**Figure 1:**
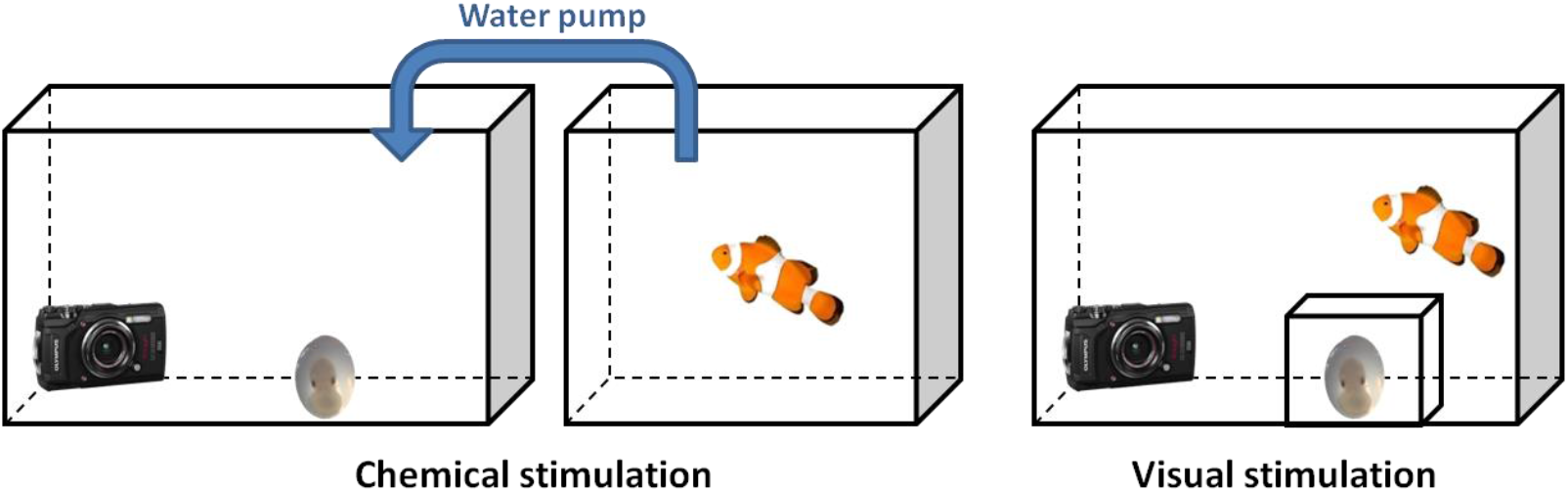
Schematic representation of the experimental device used. The cuttlefish egg was placed on the bottom in the center of the tank. The camera is positioned in front of the embryo in order to record its responses. Left part; Chemical stimulation test: fish were in another tank connected to the experimental tank with a water pump. Right part; Visual stimulation test: fish were placed into the embryos’ tank but the embryo was enclosed in a transparent glass container.

For the chemical stimulation, the fish aquarium (predator or non-predator) was placed next to the embryo’s tank and connected with a water pump. During the stimulation phase, the pump was turned on so that the fish odour (predator or non predator) arrived just next to the embryo (80 mL/min). For the ink condition, a sample of 3ml of ink (cf. above) was added to 150 mL of blank seawater and mixed until the solution came more or less translucent (despite the mixing, a light grey colouration can persist). A sample of 3 mL of this solution was then presented to the embryo; since the tank is dark, the embryo could probably not see the ink (see Boal & Golden 1999). For the visual stimulation, the fish was directly placed in the embryo’s tank. The embryo was placed in a transparent glass container (6×4×4cm) to protect it from the fish and avoid a chemical exposure to the predator odour. For the ink condition, 3ml of black ink was presented next to the embryo.

### 1) Innate recognition test

#### a) Innate chemical recognition test

Experimental stimuli used were:

1. Blank seawater (“**C**”, control condition); n=9
2. Odour of Clownfish (“**NP**”, non-predator condition); n=8
3. Odour of Puffer fed with shrimp (“**P_shrimp_**”, predator condition); n=6
4. Odour of Puffer fed with cuttlefish embryos (“**P_embryo_**”, predator condition); n=6
5. Cuttlefish ink (“**I**”, ink condition); n=17.

#### b) Innate visual recognition test

Experimental conditions were:

1. Clownfish (“**NP**”, non-predator condition); n=8
2. Puffer (“**P**”, predator condition); n=10
3. Black cuttlefish ink (“**I**”, ink condition); n=12.

The last minute of the acclimation and the first minute under experimental condition (stimulation time) were recorded. Data collection was carried out by counting manually the ventilation rate (VR) each minute. Preliminary studies showed that embryos respond immediately when they are exposed to stimulation: during acclimation VR did not change but it change during the stimulation phase. The observer was blind to the treatments.

### 2) Learned recognition test

#### a) Conditioning phase

A classical conditioning procedure has been used. The Clownfish (NP; non-predator) was used as a conditional stimulus (CS) and the cuttlefish ink (I) was used as an unconditional stimulus (US). Embryo was exposed to the CS coupled with the US once a day for 30 min during 4 days. The stimuli used in this experiment were obtained according to the same procedure as in the innate recognition tests described above.

Two groups were tested for the chemical recognition:

1. The first group, experimental group “**NP+I**” (n=12), included embryos exposed to a clownfish odour (Non-Predator) paired with cuttlefish ink odour.
2. The second group, control group “**NP**” (n=12), included embryos exposed to a clownfish odour alone (Non-Predator).

Two groups were tested for the visual recognition:

1. The first group, experimental group “**NP+I**” (n=10), included embryos exposed to a clownfish (Non-Predator) coupled/paired with cuttlefish ink clouds.
2. The second group, control group “**NP**” (n=12), included embryos exposed to a clownfish alone (Non-Predator).

#### b) Testing phase

On the 5^th^ day, all embryos were tested with the odour or the sight of clownfish alone. Data collection was carried out by counting manually the VR one minute before and after the stimulation phase. The observer was blind to the treatments.

### 3) Statistical analyses

Given the sample size, nonparametric statistical methods were used to analyse data. Mean ventilation rates during acclimation period and stimulation phase were compared using a Wilcoxon test (R©3.2.0). The α level for all analyses was 0.05. For the graphical representation, each histogram bar represents the index calculated as follows:

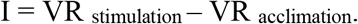

This index shows whether the RV increases or decreases as a result of stimulation (positive values mean that the VR increases after the stimulation; negative values mean that the VR decreases).

### 4) Ethical note

All animals (fishes and cuttlefish) and the entire protocol were approved by the National Tsing Hua University Institutional Animal Care and Use Committee (IACUC Protocol No. 10510). Throughout the protocol, we followed the published guidelines for the care and welfare of cephalopods to avoid stress in test animals (Fiorito et al., 2015).

## Results

### 1) Innate recognition

The ventilation rate (VR) of each embryo was measured before stimulation (VR_acclimation_) and after stimulation (VR_stimulation_). By using the index I (I = VR_stimulation_ - VR_acclimation_), we can then see whether the RV decreases or increases as a result of this stimulation.

#### a) Chemical

Embryos’ VR did not change after exposure to blank seawater (“C” group); to non-predator odour (“NP” group) and to odour of the predator fed by shrimp (“P_shrimp_” group) (Figure 2: C group: Z=0.00; p>0.999; NP: Z=−0.57; p=0.574; P_shrimp_: Z=−1.134; p=0.257). Embryos’ VR decreased after exposure to the odour of predator fed by cuttlefish embryo (“P_embryo_” group) and to ink odour (“I”group) (Figure 2: P_embryo_: Z=−2.041; p=0.041; I: Z=−2.650; p=0.008).

**Figure 2:**
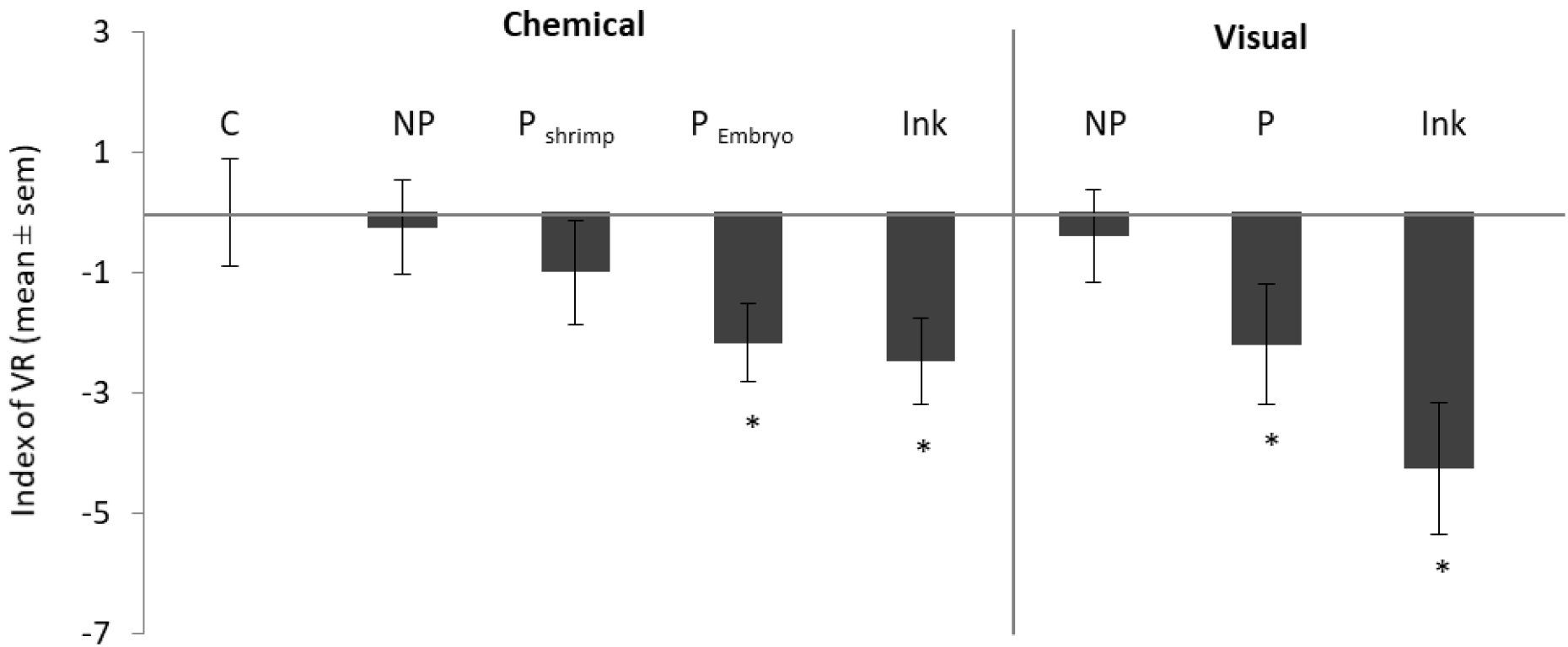
Index of ventilation rate (VR) of embryos exposed to blank seawater (C); non-predator (NP); predator (P - with P_shrimp_ = predator group fed with shrimp (puffer) and P_embryo_ = predator group fed with cuttlefish embryo; cuttlefish ink (I). Wilcoxon test: *: p<0.05.

#### b) Visual

Embryos’ VR did not change after exposure to non-predator (“NP” group) (Figure 2: NP: Z=−0.537; p=0.590). Embryos’ VR decreased after exposure to predator (“P” group) and to ink (“**I**” group) (Figure 2: P: Z=−2.025; p=0.042; I: Z=−2.83; p=0.0047).

### 2) Learned recognition

#### a) Chemical

After 4 days of repeated exposure to clownfish odour paired with ink odour, VR significantly decreased when embryos were exposed to clownfish odour alone the 5^th^ day (Figure 3: NP+I: Z=−2.157; p=0.031). On the contrary, after 4 days of repeated exposure to clownfish odour alone, VR did not change if embryos were exposed to clownfish odour alone the 5^th^ day (Figure 3: NP: Z=−0.303; p=0.762).

**Figure 3:**
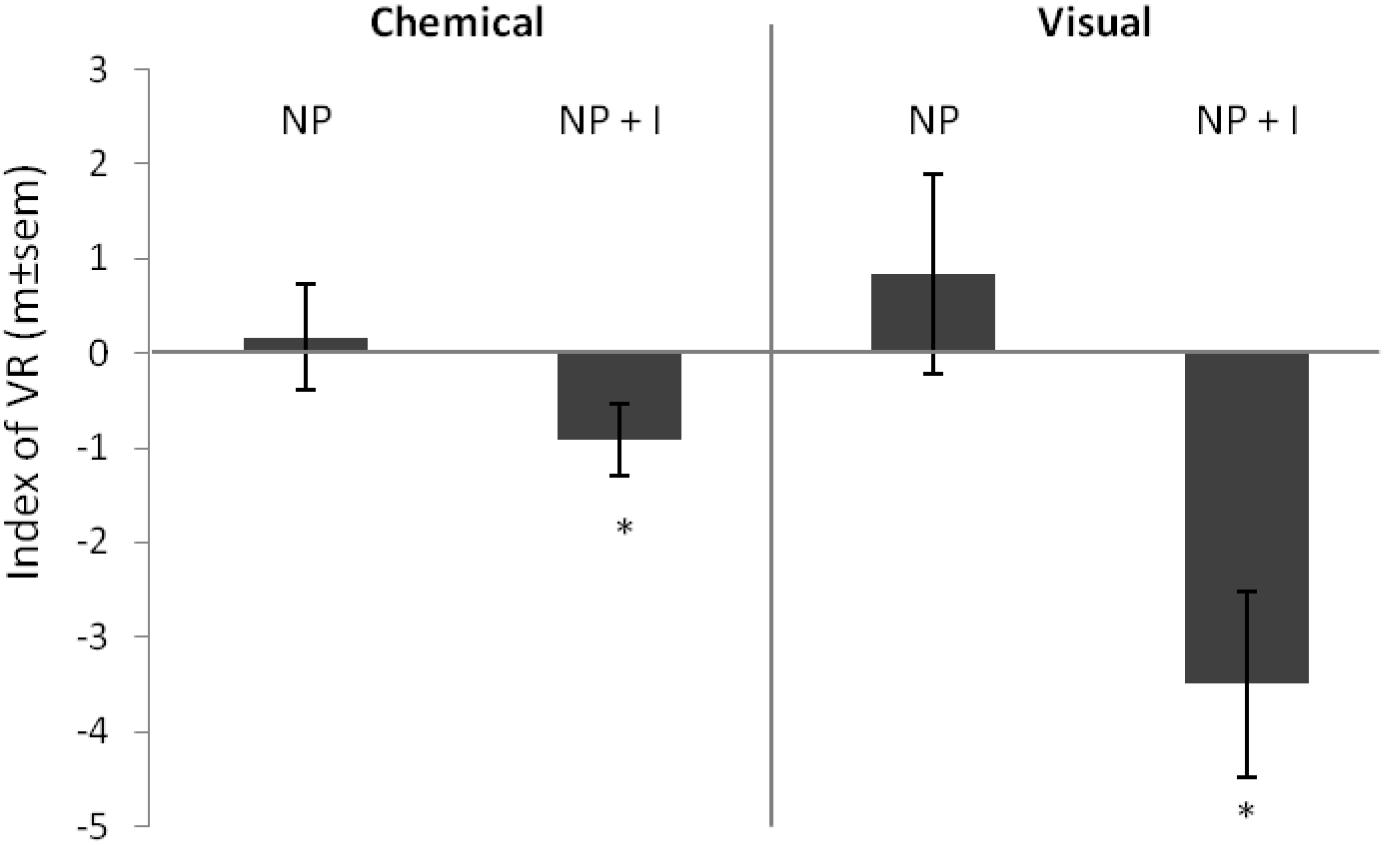
Index of ventilation rate (VR) of embryos after associative learning protocol. Associative learning condition: NP=non-predator only during 4 days (clownfish, control group); NP+I=non-predator coupled with cuttlefish ink during 4 days. Wilcoxon test: ns: p≥0.05; *: p<0.05.

#### b) Visual

After 4 days of repeated exposure to clownfish paired with ink, VR significantly decreased when embryos were exposed to clownfish alone the 5^th^ day (Figure 3: NP+I: Z=−2.395; p=0.017). Conversely, after 4 days of repeated exposure to clownfish odour alone, VR did not change if embryos were exposed to clownfish odour alone the 5^th^ day (Figure 3: Z=0.714-; p=0.475).

## Discussion

In the first part of this study, we investigated whether embryos can innately recognize predator by using visual or olfactory cues. We showed that ventilation rate (VR) of *Sepia pharaonis* embryos significantly decreased when they were exposed to potential predators and ink. Changes in physiological parameters such as heart rate or VR often demonstrate perception (Colombelli-Négrel, Hauber, & Kleindorfer, 2014; Oulton, Haviland, & Brown, 2013) or attention abilities (Porges & Raskin, 1969; Richards & Casey, 1991), notably when the animal is in a dangerous situation. The VR often increases to prepare an individual for flight to avoid a predator (Misslin, 2003). However, predator detection through visual or chemical stimulus could also induce a “freezing-like” behaviour (Misslin, 2003) along with a decrease of VR. In mammals, freezing is considered to be a fear response related to a harmful stimulus, characterized by immobility and changes in physiological parameters, such as cardiac and VR, and may enhance a prey’s survival facing predation. In cuttlefish, few studies focused on changes in the VR during the stimulation. In these studies, change of VR reflect visual or chemical perception abilities (Boal & Golden, 1999; Boal & Ni, 1996; King & Adamo, 2006). Indeed, juvenile cuttlefish became motionless (behavioural freezing), hyperinflated their mantle, and decreased their VR and heart rate upon presentation of the sudden visual stimulus (rapidly approaching bird cut-out) (King & Adamo, 2006). Likewise, adult cuttlefish decreased breathing was associated with freezing-like response and this freezing response seems adaptive since it could reduce the risk of being detected by movement, and also reduces the bioelectric field, which may prevent attacks by sharks (Bedore, Kajiura, & Johnsen, 2015). *Sepia pharaonis* eggs are totally transparent; consequently, reducing movements associated with reducing breath and hence general activity inside the egg may reduce the probably for the embryo to be detected by predators and increase their survival chances. VR is also a sensitive indicator of fish physiological responses to stress (Barreto et al., 2003). In their study, Barreto et al. measured the VR of the Nile tilapia (*Oreochromis niloticus*) before and after the presentation of three stimuli: an aquarium with a harmless fish or a predator or water (control). Nile tilapia VR increased significantly in the group visually exposed to a predator compared with the other two which indicates a recognition ability (Barreto et al., 2003).

As in vertebrate species, we showed that cuttlefish embryos innately respond to chemical cues from predators but not from non-predators. Indeed, our study shows that embryos respond differently to puffer fed with frozen shrimp (less harmful) and puffer fed with cuttlefish embryos (harmful). The VR significantly increased only when embryos were exposed to the latter. This result suggests that embryos do not respond to the fish odour itself but rather to the degree of dangerousness of the predator based on its diet. This specific recognition is in accordance with the results of a study on the clownfish *Amphiprion percula*, in which larvae anemonefish (*Amphiprion percula*) showed indifference to chemosensory cues from non-piscivorous fishes fed their usual diet, but significantly avoided chemical cues from piscivorous and non-piscivorous fishes fed a diet containing fish product (Dixson, Pratchett, & Munday, 2012).

One of the most noteworthy results of the present study is that predator recognition is not only based on chemical cues but also on predatory visual information. VR decreased when embryos were exposed to the puffer but not to the clownfish. This change of VR cannot be attributed to a lack of oxygen; the egg is enclosed in a box and first, we did not observe any change of VR when embryos were exposed to the non-predator and second, because a lack of oxygen would have rather caused an increase in VR (Randall, & Shelton, 1963). Which visual predatory cues embryos have used is a question that remains unanswered. Nevertheless some hypotheses can be proposed. First, since the size of the fish has been controlled, the recognition can be based on their behaviour. Indeed, the fish behaviour in the experimental tank was different between the two species. The puffer fishes have been trained to prey on the eggs and thus cuttlefish embryos are the basis of their diet. As a consequence, during the the exposure, the puffer spent most of the time close to the glass box and directed several attacks towards the egg (personal observation). On the contrary, the clown fish stayed away from the egg mostly close to the side of the aquarium. Consequently the level of threat is higher with the puffer than with the clownfish. This observation is in accordance with other ones on the same model ; juveniles cuttlefish display secondary behaviour (deimatic pattern and inking) when the puffer fish is close to them (Lee, Darmaillacq, Dickel, & Chiao, submitted). Second, a morphometric analysis of 20 different facial features of reef fishes was carried out in order to assess cues which could serve for predator recognition and showed that the shape of the fish’s mouth and the distance between the eyes and the mouth could be different between a carnivorous and an herbivorous fish (Karplus & Algom, 1981). This morphological criterion may be sufficient for good visual recognition of predator.

Our study highlights that embryos innately respond to the sight of a ink cloud as well as to the ink odour at a very low concentration as a warning signal. Again, this response is adaptive because it decreases probability to be detected by predators that potentially attacked eggs or hatchlings in the vicinity of eggs. In fish and amphibian species, young individuals innately respond to chemical alarm cues (pheromones) released by injured conspecific. In cephalopod, threatened individuals jet clouds of black ink. The cuttlefish ink can be a relevant warning signal (Derby, 2014). It is composed of secretions from two glands: (1) the gland of the ink bag that produces a black ink containing melanin; (2) the gland in the funnel that produces mucus. The cuttlefish ink is composed of melanin but also catecholamines, DOPA and dopamine (which are monoamines derived from tyrosine), amino acids such as taurine but also certain metals such as cadmium, copper and lead (Derby, 2014; Madaras et al., 2010; Prota et al., 1981). The ink of cephalopods would then have a role of defense against predators in two ways: (1) ink as a direct deterrent of predators (interspecific effects) and (2) Ink as an alarm cue for conspecifics (intraspecific effect) (Derby, 2014; Hanlon & Messenger, 2018). This second type of defense acts indirectly against predators, because it signals a danger to conspecifics. In *Loligo opalescens* squid ink can cause inking and camouflage (Gilly & Lucero, 1992; Lucero, Farrington, & Gilly, 1994). Also, dopamine at biologically relevant concentrations is sufficient to cause ink ejection (Gilly & Lucero, 1992; Lucero et al., 1994).

In the second part of the study, we showed that embryos can learn to recognize a stimulus as being harmful when it is paired with ink. Indeed, we showed that VR of cuttlefish embryo decreased significantly to the presentation of a clownfish alone (through sight or smell) when it was coupled with ink the 4 days before. It is unlikely that this group may be experiencing sensitisation. Unpublished data show that pairing the clownfish odour with ink for 2 days leads to the same results in *S. pharaonis* and also in *S. officinalis* after only one pairing. We also showed that embryos exposed to cuttlefish ink once a day for four days do not respond to neutral odour (cinnamon; personal observation) afterwards. In the present study, it is then likely that embryos are able to learn to recognize a new predator by associative learning. Associative learning, defined as a learned link between two events or between a behaviour and its consequences (Bouton, 2007), has been shown in cuttlefish (adults and juveniles) and other cephalopod including octopuses (O’Brien, Mezrai, Darmaillacq, & Dickel, 2017; Wells, 1968; Young, 1961). Cuttlefish (*S. officinalis*) can learn the visual characteristics of prey while inside the eggs by a mere exposure, thus non associative learning because juveniles’ spontaneous food preferences are altered after an embryonic crab exposure (Darmaillacq et al., 2008, 2017; Guibé et al., 2012)). The present study shows direct evidence that cuttlefish embryos can also learn through classical conditioning. This learning capability is adaptive given that it allows cuttlefish to learn information about its future environment while it is safe inside the egg case, and hence improves the survival chances of juveniles after hatching. These results are in accordance with studies on tadpoles and invertebrate larvae in which embryos can also learn about new predators when they are paired with alarm cues (mosquitoes: Ferrari et al., 2008; damselfly: Wisenden et al., 1997). Predation is a constant threat faced by most prey individuals. Learning about predation before hatchling is a great advantage for the survival of young animals, especially if they develop without direct parental care.

To conclude, being able to detect and learn about predators is highly beneficial for the embryo while still protected by the egg case. In a changing environment, these prenatal learning abilities are important in case of new predators (e.g. invasive species) or in case of predator diet changes. Indeed, in fish, the flexibility of feeding behaviour is an important adaptive trait because most natural environments change spatially and temporally (Dill, 1983; Vehanen, 2003; Wright, Eberhard, Hobson, Avery, & Russello, 2010). Developing in a transparent egg allows using visual information in addition to chemosensory ones. Last, in this study, cuttlefish embryos are able to learn in 4 days, but unpublished experiments showed that learning can be faster: 2 days in *Sepia pharaonis*) and 1 day in *Sepia officinalis;* this allows embryo to learn until the last days before hatching.

## Acknowledgements

This work was supported by a grant from Campus France (Orchid project 2017-2018) and by the Agence Nationale de la Recherche (ANR PReSTO’Cog 13-BSV7-0002). The authors gratefully acknowledge Jane Martin for editing the English in this manuscript.

